# A Multiplexed DNA FISH strategy for Assessing Genome Architecture in *C. elegans*

**DOI:** 10.1101/397471

**Authors:** Brandon Fields, Son C. Nguyen, Guy Nir, Scott Kennedy

## Abstract

Eukaryotic DNA is highly organized within nuclei and this genomic organization is important for genome function. Fluorescent *in situ* hybridization (FISH) approaches allow the 3D architecture of genomes to be visualized. Scalable FISH technologies, which can be applied to whole animals, are needed to help unravel how genomic architecture regulates, or is regulated by, development, growth, reproduction, and aging. Here, we describe a multiplexed DNA FISH Oligopaint library that targets the entire *C. elegans* genome at chromosome, three megabase, and 500 kb scales. We describe a hybridization strategy that provides flexibility to DNA FISH experiments by coupling a single primary probe synthesis reaction to dye conjugated detection oligos via bridge oligos, eliminating the time and cost typically associated with labeling probe sets for individual DNA FISH experiments. The approach allows visualization of genome organization at varying scales in all/most cells across all stages of development in an intact animal model system.

## Introduction

Eukaryotic genomes are non-randomly organized within mitotic and interphase nuclei. The basic unit of genome organization is the nucleosome, which assemble into higher order structures whose biogenesis, maintenance, regulation, and purpose are poorly understood (Bonev and Cavalli 2016; Meaburn 2016; Yu and Ren 2017; Bickmore 2013). DNA fluorescent *in situ* hybridization (FISH) technologies and chromatin conformation capture techniques allow 3D architectures of genomes to be assessed (Dekker et al. 2002; Lieberman-Aiden et al. 2009; Bauman et al. 1980; Beliveau et al. 2012; Dekker, Marti-Renom, and Mirny 2013; Bienko et al. 2013). Studies using these technologies have begun to reveal how DNA is organized within nuclei. For instance, chromatin capture experiments indicate that many eukaryotic genomes are assembled into megabase-sized structures termed topologically associated domains (TADs). TADs, and larger organization units termed compartments, are thought to allow subregions of chromosomes to share and integrate long-range transcriptional regulatory signals (Dixon et al. 2012; Dekker and Heard 2015; Dekker and Mirny 2016; Vernimmen and Bickmore 2015; Lieberman-Aiden et al. 2009). Additionally, DNA FISH and chromatin immunoprecipitation (ChIP) experiments have shown that the position of genes within nuclei is often not random: active genes tend to localize near nuclear pores and/or the nuclear interior while inactive genes tend to localize to the nuclear periphery, distant from nuclear pores (Pickersgill et al. 2006; Casolari et al. 2004; van Steensel and Belmont 2017; Gonzalez-Sandoval and Gasser 2016; Lemaître and Bickmore 2015). Finally, DNA FISH experiments have shown that individual chromosomes tend to occupy distinct non-overlapping regions, even in interphase nuclei (termed chromosome territories) (Cremer and Cremer 2010; Bolzer et al. 2005). Many questions concerning the large-scale architecture of genomes remain unanswered, including the following: how the various aspects of genome architecture, such as gene position, TADs, or territories differ in different cell types or across developmental time, and how such changes relate to gene expression. Technologies that enable rapid and flexible analysis of genome organization in an intact animal would allow such questions to be addressed.

DNA FISH experiments are in a large part limited by the cost and time associated with producing fluorescent probe sets. Oligopaint technology has made probe production faster and cheaper (Beliveau et al. 2012). Oligopaints takes advantage of massively parallel DNA synthesis technologies to create libraries containing hundreds of thousands of individual DNA oligos each comprised of a short (42 bp) DNA sequence that hybridizes to a genome, as well as additional “barcode” sequences important for other aspects of library function. For instance, barcode sequences allow oligos to be repeatedly amplified from an Oligopaint library using PCR and T7 polymerase based approaches, thus providing a virtually inexhaustible supply of oligos for DNA FISH experiments (Beliveau et al. 2012; Chen et al. 2015; Murgha, Rouillard, and Gulari 2014). Also, while traditional FISH probes utilize fluorescent dyes covalently attached to FISH probes, barcode sequence obviate the need for labeling new probe sets prior to each DNA FISH experiment by allowing pre-labeled detection oligos to be used to detect Oligopaint oligos (Beliveau et al. 2015). Here, we describe a modified Oligopaint strategy that empowers rapid, flexible, and inexpensive DNA FISH. We report methods for using this library to simultaneously visualize all six *C. elegans* chromosomes, as well as three megabase and 500 kilobase subregions of these chromosomes, in all/most cells of *C. elegans* across all stages of development.

## Results

### *C. elegans* Oligopaint library design

The Oligopaints bioinformatics pipeline was used to identify 42 bp DNA sequences in the *C. elegans* genome (Ce 10) that 1) uniquely map to the genome, 2) exhibit similar melting temperatures and similar GC content, 3) lack repetitive stretches, and 4) lack predicted secondary structures (Beliveau et al. 2012). Detailed parameters for this analysis are described in materials and methods. We used the results of this bioinformatic search to generate an Oligopaint library that contained approximately 2 probes/kb of genomic sequence evenly spaced across each *C. elegans* chromosome (Fig, 1a and Table S1). In total, the library consisted of 170,594 oligos (termed primary Oligopaint oligos), which each contain 42 bp of unique genomic sequence flanked by barcode sequences that allow for DNA FISH targeting each of the six *C. elegans* chromosomes, as well as three megabase, or 500kb subregions of these chromosomes (Fig. 1b). Bridge oligos (also see Guy Nir, bioRxiv 2018) were designed that base pair with barcode sequences engineered into primary probes as well as base pair with dye-conjugated detection oligos (Fig. 1c). Detection oligos were designed that base pair with bridge oligos and are conjugated to three fluorophores (Alexa 488, Cy3, and Alexa 647) (Fig. 1c). Bridge oligos serve as an intermediate oligo that hybridizes to the primary probe and provide a docking site for detection probes. Thus, bridge oligos provide versatility (and cost savings) to DNA FISH experiments as these oligos allow any primary probe set to be coupled to any detection probe set with minimal additional cost. Bridge oligos also allow for more than one fluorophore to be targeted to primary probes, which expands the number of objects that can visualized with a standard three channel microscopy system (see 6 chromosome FISH experiments below). By using detection oligos with fluorophores on both 3’ and 5’ termini, with two detection oligos per bridge oligo, and using bridge oligos that target the 5’ and 3’ barcode sequence of primary probes, it is possible to have each primary oligo recognized by eight fluorophores. To visualize *C. elegans* chromosomes, unlabeled primary probes are first PCR amplified from the Oligopaint library as described in (Chen et al. 2015; Murgha, Rouillard, and Gulari 2014; Beliveau et al. 2012)) (also see materials and methods). Second, primary Oligopaint oligos are hybridized to fixed samples of *C. elegans* overnight (see below and materials and methods). Third, samples are hybridized with a mixture of bridge oligos and dye-conjugated detection oligos for 3 hours the following day (Fig. 1c). Together, the three step strategy described above allows many DNA FISH experiments to be conducted fairly cheaply after a single primary probe synthesis step.

**Figure 1.**
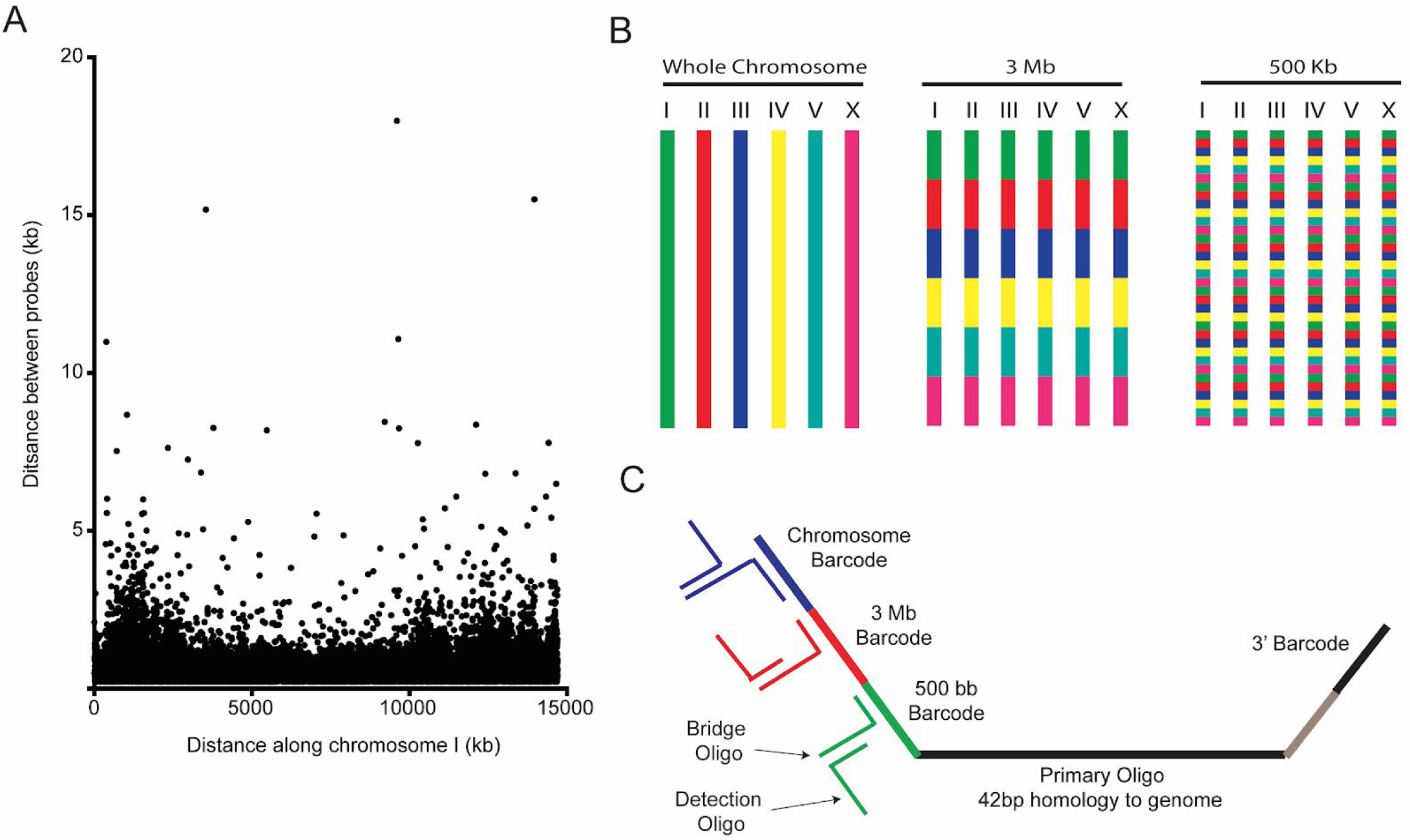
A highly multiplexed oligo library for *C. elegans* Oligopaint. **(A)** The distribution of Oligopaint probes across chromosome I is shown. Probe distribution is similar for other chromosomes (see Table S1). **(B)** Oligopaint library allows primary oligos, which are specific to any chromosomes, 3Mb, or 500Kb region within any chromosome, to be specifically amplified. Primary probes are PCR amplified from Oligopaint library and produced as described in materials and methods. Bridge and detection probes allow the indicated chromosomal regions to be visualized. **(C)** Primary probes consist of barcode sequences appended to 42 bp sequences that hybridizes uniquely to *C. elegans* genome. Total length of each oligo is 150 bp. Barcode sequences allow each primary probe to amplified as part of a pool of primary probes that target a chromosome (chromosome barcode), 3Mb subsection of chromosome (3 Mb barcode), or 500 kb subsection of chromosome (500 kb barcode). Bridge oligos and Detection oligos (arrows) are used to recognize and illuminate primary probes. Note: primary oligos contain an additional barcode not used in this work (brown). The barcode is specific to each 500 kb segment and could be used to increase detection efficiencies of 500 kb DNA FISH by allowing an addition detection oligo to be incorporated during the detection phase of DNA FISH.

### *C. elegans* Oligopaint staining is robust and specific

DNA FISH in *C. elegans* is typically done on dissected tissue. We developed a fixation and hybridization protocol that allowed for efficient DNA FISH on intact *C. elegans*. As part of this protocol, hybridization steps are conducted in microcentrifuge tubes allowing large numbers of animals to be simultaneous assayed by FISH. A detailed description of this fixation and hybridization protocol can be found in materials and methods. To test our *C. elegans* Oligopaint library, we amplified a primary probe set targeting chromosome II (27,360 unique probes) and asked if this probe set was able to specifically label chromosome II. The behavior and morphology of chromosomes in the *C. elegans* germline are well-established (Albertson, Rose, and Villeneuve 2011). For instance, homologous chromosomes pair at the pachytene stage of Meiosis I at a defined region of the germline (termed pachytene region). Oocytes are arrested in diakinesis of meiosis I and chromosomes are highly condensed with homologs connected via a single chiasmata (termed bivalents) (Villeneuve 1994). Mature sperm harbor highly condensed chromosomes and are haploid. To address specificity, we imaged germlines of animals subjected to our DNA FISH approach targeting chromosome II. This analysis detected the expected chromosomal structures in pachytene germ cells, oocytes, and sperm; fluorescent staining was observed on a single bivalent in oocytes and in one region of the nucleus in sperm and pachytene germ cells (Fig. 2). The data show that the *C. elegans* Oligopaint library is specific. To quantify the efficiency of our method, we first measured the % of whole animals stained by chromosome II DNA FISH. Staining could be observed in 1085/1303 (83%) of larval stage animals, 317/326 (97%) of adult animals, and 50% of the embryos housed within uteri of adult animals (Fig. S1). Note, an alternative protocol that allows for greater efficiency in embryos, but not intact animals, is described in materials and methods. We next measured the % of nuclei within a given animal that were stained by chromosome II DNA FISH. We randomly chose DAPI-stained nuclei (from animals that showed staining) and asked if these nuclei were positive for chromosome II DNA FISH signals. Out of 50 randomly chosen somatic nuclei 50/50 had DNA FISH signal.

**Figure 2.**
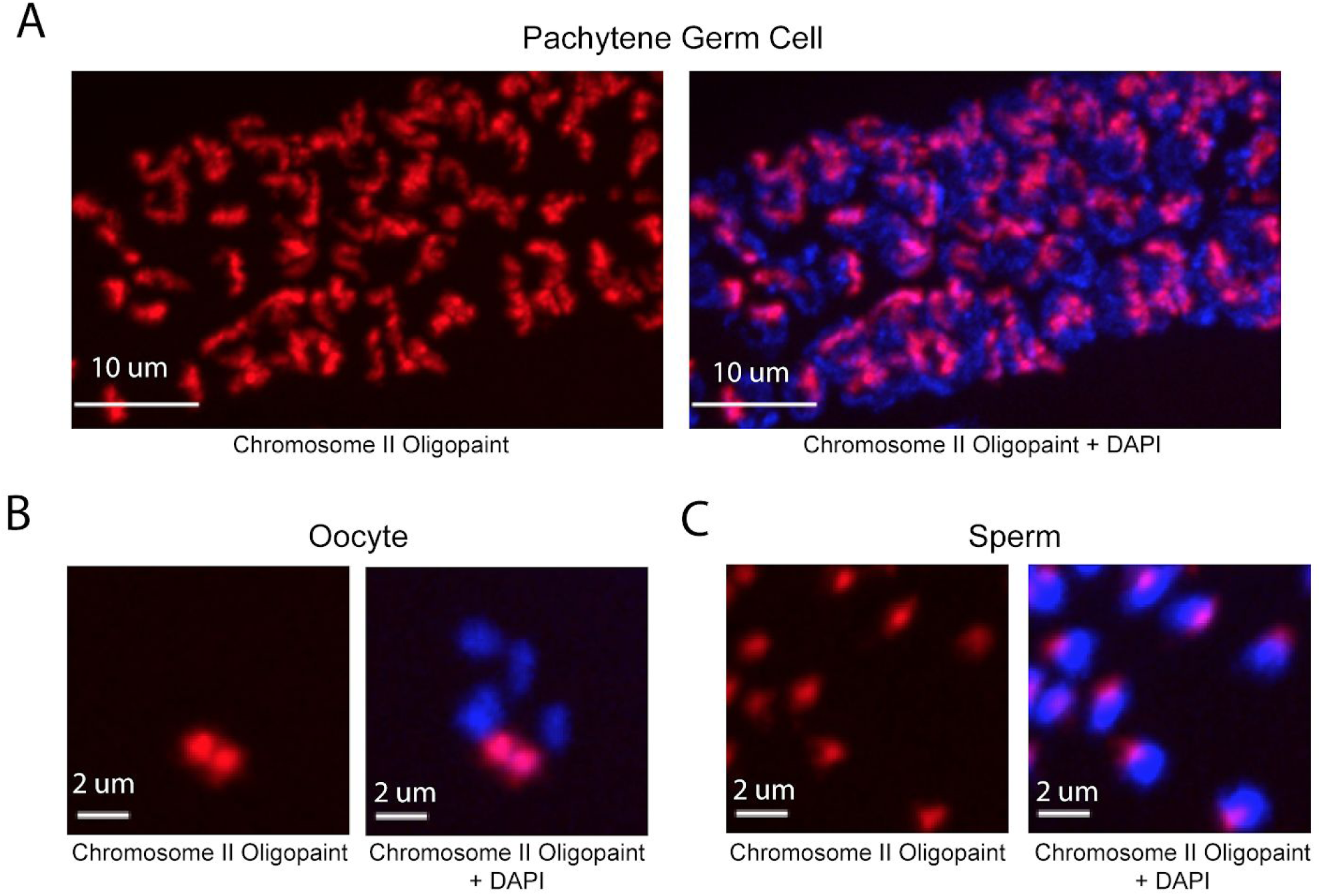
Whole chromosome Oligopaint in *C. elegans* is specific. (A-C) Adult *C. elegans* were fixed as described in materials and methods. Fixed animals were subjected to three step hybridization to detect Chromosome 2 (red). Animals were co-stained with DAPI (blue). **(A)** 3D maximum projection of the pachytene region of an adult *C. elegans* germline is shown. **(B)** 3D maximum projection of an oocyte. A single bivalent is stained with chromosome II Oligopaints. **(C)** 3D maximum projection of sperm. Scale bars for images are indicated.

Likewise, 50/50 germline nuclei were positive for DNA FISH signals. We conclude that our library and hybridization strategy allows for robust and specific labeling of a whole chromosome in many cell types and many developmental stages simultaneously in large numbers of animals. It is possible that DNA FISH signals in every cell and at every stage of development can be visualized with this approach.

### Simultaneous visualization of all six *C. elegans* chromosomes

To detect all six *C. elegans* chromosomes simultaneously, we amplified primary probe sets targeting all six *C. elegans* chromosomes and hybridized these primary probes to fixed adult *C. elegans*. We then used bridge oligos to couple primary probe sets to combinations of detection oligos labeled with Alexa 488, Cy3, and Alexa 647 (Fig. 3a). Differing combinations of these three dyes on each individual chromosomes should produce six distinct colors (Fig. 3a). We imaged oocytes in these animals and detected six bivalents that were each labeled a distinct color (Fig. 3b). DNA FISH staining was robust: 50/50 randomly chosen DAPI positive oocyte nuclei were stained positive for all six colors. In pachytene stage adult germ cells, *C. elegans* chromosomes are paired, condensed, and localized near the nuclear periphery (Albertson, Rose, and Villeneuve 2011). DNA FISH illuminated six regions of distinct colors concentrated near the nuclear periphery in pachytene germ cells (Fig. 3c). Six chromosome FISH staining was also successful in somatic nuclei. Six distinct colors were often distinguishable in the nuclei of intestinal and hypodermal nuclei, as well as ventral cord nuclei whose small size and positioning within the animal were indicative of ventral cord neurons (Fig. 3d-f and Movie 1). These data show that our DNA FISH approach is capable of labeling all six *C. elegans* chromosomes simultaneously across different cell types of the intact animal. Incidentally, the data also show that, like interphase chromosomes in other eukaryotes, *C. elegans* chromosomes occupy largely distinct territories within interphase nuclei and that chromosome territories can persist in post-mitotic cells.

**Figure 3.**
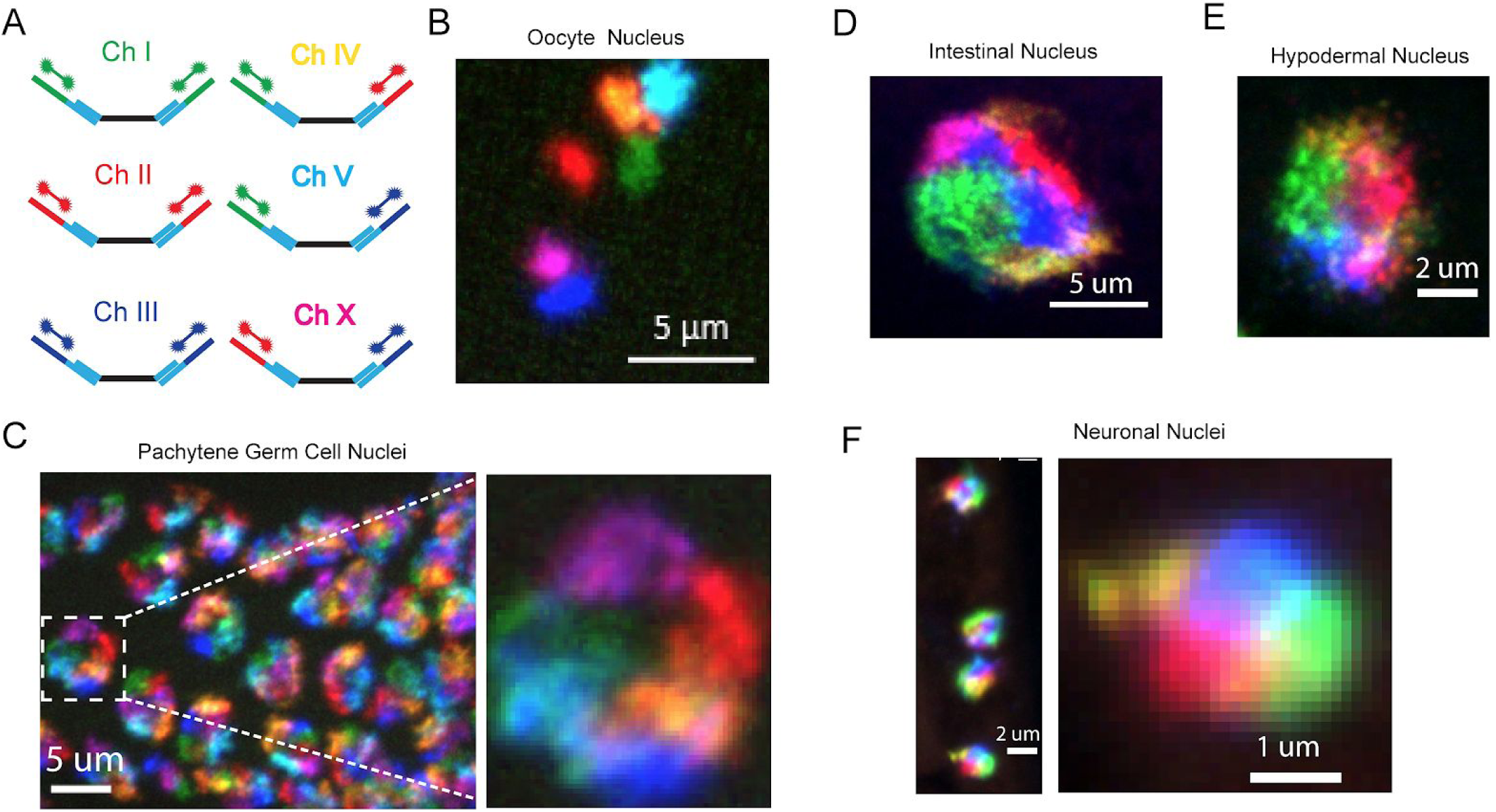
Simultaneous visualization of all six *C. elegans* chromosomes. **(A)** Strategy to detect six chromosomes is shown. Detection probes labeled with Alexa488 (Green), Cy3 (Red), and Alexa647 (Blue), or combinations of these three fluorophores were used to label each of the six *C. elegans* chromosome a different color. **(B-F)** Adult *C. elegans* were fixed and subjected to three step hybridization to detect Chromosomes 1, 2, 3, 4, 5, and X. **(B)** 3D maximum projection of an oocyte. Each bivalent is labeled a different color. **(C)** 3D maximum projection of the pachytene region of an adult germline. A magnification of one of these nuclei is shown to the right. **(D)** 3D maximum projections of an intestinal nucleus, **(E)** hypodermal nucleus, and **(F)** a nuclei whose size and position within the animal suggest the cell is a ventral cord neuron.

### Detection of 3Mb and 500kb chromosomal subregions

We designed our Oligopaint library to include 3 Mb and 500 Kb barcode sequences that should permit visualization of chromosomal subregions (Fig. 1c). To test this aspect of our library, we amplified Oligopaint oligos targeting chromosome I and hybridized these probes to adult *C. elegans*. We then used bridge oligos that recognized all Chromosome I Oligopaint oligos (∼13 Mb), a 3 Mb subregion of chromosome I (0-3 Mb), or a 500 kb subregion of this 3 Mb region (1.0-1.5 Mb). Detection oligos coupled to Alexa 488, Cy3, and Alexa 647 were used to illuminate each genomic region, respectively. We imaged pachytene germ cells and, as expected, observed a single contiguous DNA FISH signal after chromosome I DNA FISH (Fig. 4a). 3 Mb DNA FISH illuminated a subregion of the chromosome I signal and 500 kb DNA FISH illuminated a subregion of this 3 Mb signal (Fig. 4a). Staining was robust, with 50/50 randomly chosen nuclei possessing all three fluorescent signals. Similar patterns were observed when chromosome IV, a 3 Mb subregion of chromosome IV (0-3 Mb), or a 500 kb subregion of this 3 Mb region (2.5-3.0 Mb) were analyzed (Fig. 4b). We conclude that the Oligopaint library has the capability to visualize 3 Mb and 500 kb subregions of the *C. elegans* genome.

**Figure 4.**
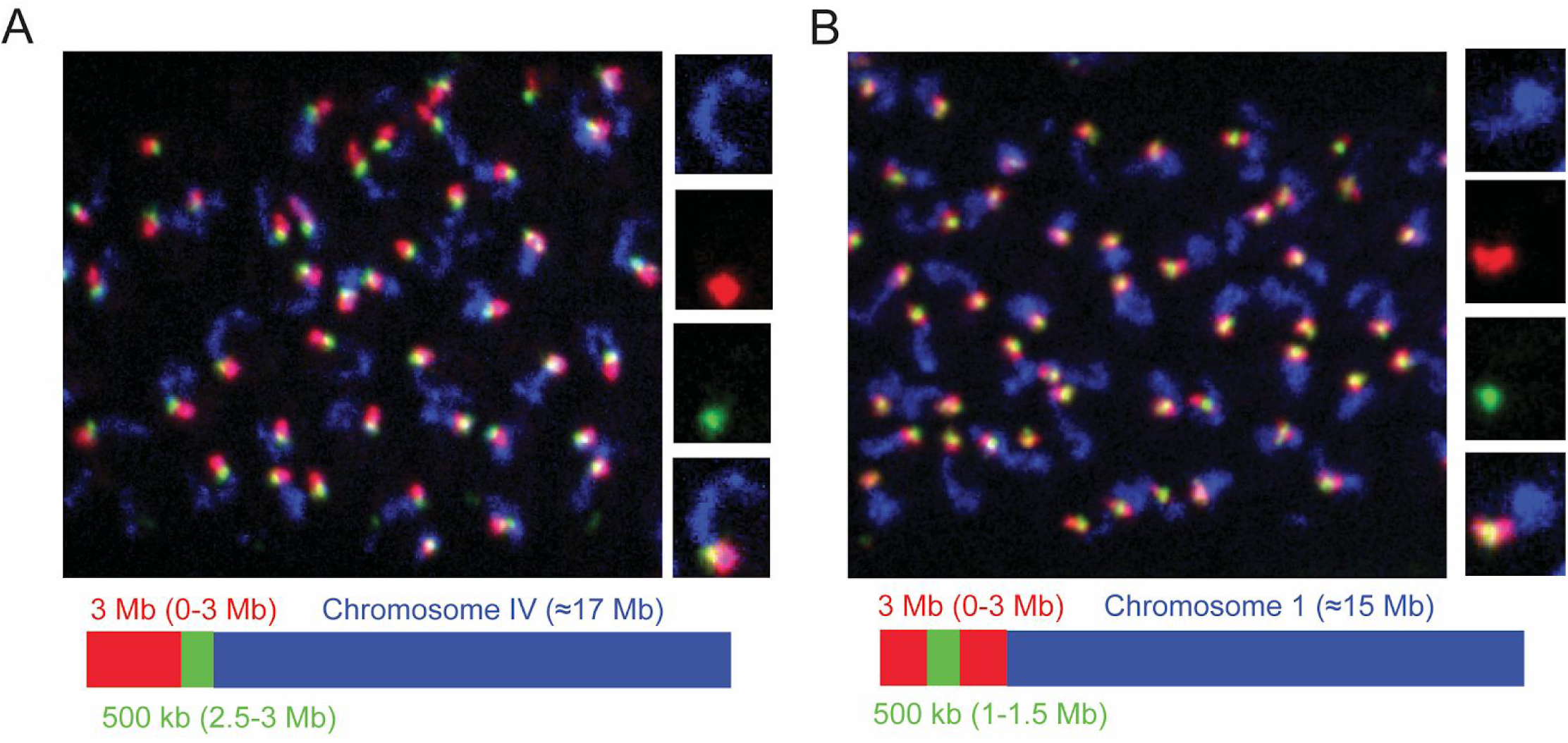
Detection of 3Mb and 500kb chromosomal subregions. Adult *C. elegans* were fixed and subjected to three step hybridization to detect Chromosome IV **(A)** or I **(B)**. **(A-B)** Top, 3D maximum project of pachytene region of adult germline. **(A)** Chromosome IV or **(B)** chromosome I, a 3MB (0-3Mb) region of these chromosomes, and a 500kb (2.5-3.0Mb) region of these chromosome were targeted with detection probes shown in blue, red, and green, respectively. Magnification of representative nuclei is shown to the right. Bottom, graphic representations of regions of chromosome IV **(A)** or chromosome I **(B)** that were stained in the experiment are shown.

### Using OligoSTORM to super resolve *C. elegans* chromosomes within whole animals

Super resolution microscopy approaches such as stochastic optical reconstruction microscopy (STORM) are able to resolve objects below the diffraction limit of light by using photoswitchable fluorophores to generate a population average of a single particle’s light emissions (Rust, Bates, and Zhuang 2006). Previous studies have coupled Oligopaints with STORM in tissue culture cells (Beliveau et al. 2015; Boettiger et al. 2016) (Guy Nir, bioRxiv 2018). We took advantage of the flexibility provided by our *C. elegans* DNA FISH protocol to ask if we could couple Oligopaints to STORM in an intact animal. We hybridized adult *C. elegans* with a primary probe set targeting chromosome IV and then used detection probes conjugated to the STORM-compatible fluorophore Alexa 647 to detect chromosome IV. We imaged oocytes with a Vutara 352 (Bruker) 3D biplane microscope and represented the resultant STORM imaging data in the form of particle density maps using pair correlation analysis (Fig. 5a, and materials and methods). This analysis revealed what appeared to be a single bivalent in each oocyte, indicating that STORM was likely successful (Fig. 5a and Movie 2). Fourier ring correlation analysis estimated our resolution to be ∼80 nm in xy, suggesting our imaging had achieved super resolution (Banterle et al. 2013). Consistent with this idea, STORM imaging revealed what appeared to be a distinct chiasmata between chromosome IV homologs, which was more difficult to resolve with standard light microscopy (Fig 5A/B and Movie 2). Interestingly, three dimensional analysis of imaging data revealed what appeared to be a high density “core” at the center of each chromosome IV homolog that was surrounded by less dense chromosomal regions (Fig. 5c and Movie 2). Because Oligopaint probes are evenly distributed across each chromosome (Fig. 1a, Table 1), STORM particle density profiles should be reflective of DNA density. Analysis of individual Z slices showed that apparent changes in DNA density gradients were not an artifact of projecting three dimensional images into two dimensional space (Movies S1). Similar density profiles were observed in somatic nuclei whose size and positioning were indicative of ventral cord neurons (Fig. 5c and Movies S2-S3). Similar density gradients were observed in both germ cell and somatic cell nuclei after chromosome III STORM (Fig. S2, Movies S4-S7). To conclude, we show that STORM can be used to visualize genomic architectures at super resolution within the context of a whole animal. Additionally, the data hint that *C. elegans* chromosomes may be arranged around central cores of densely packed DNA.

**Figure 5.**
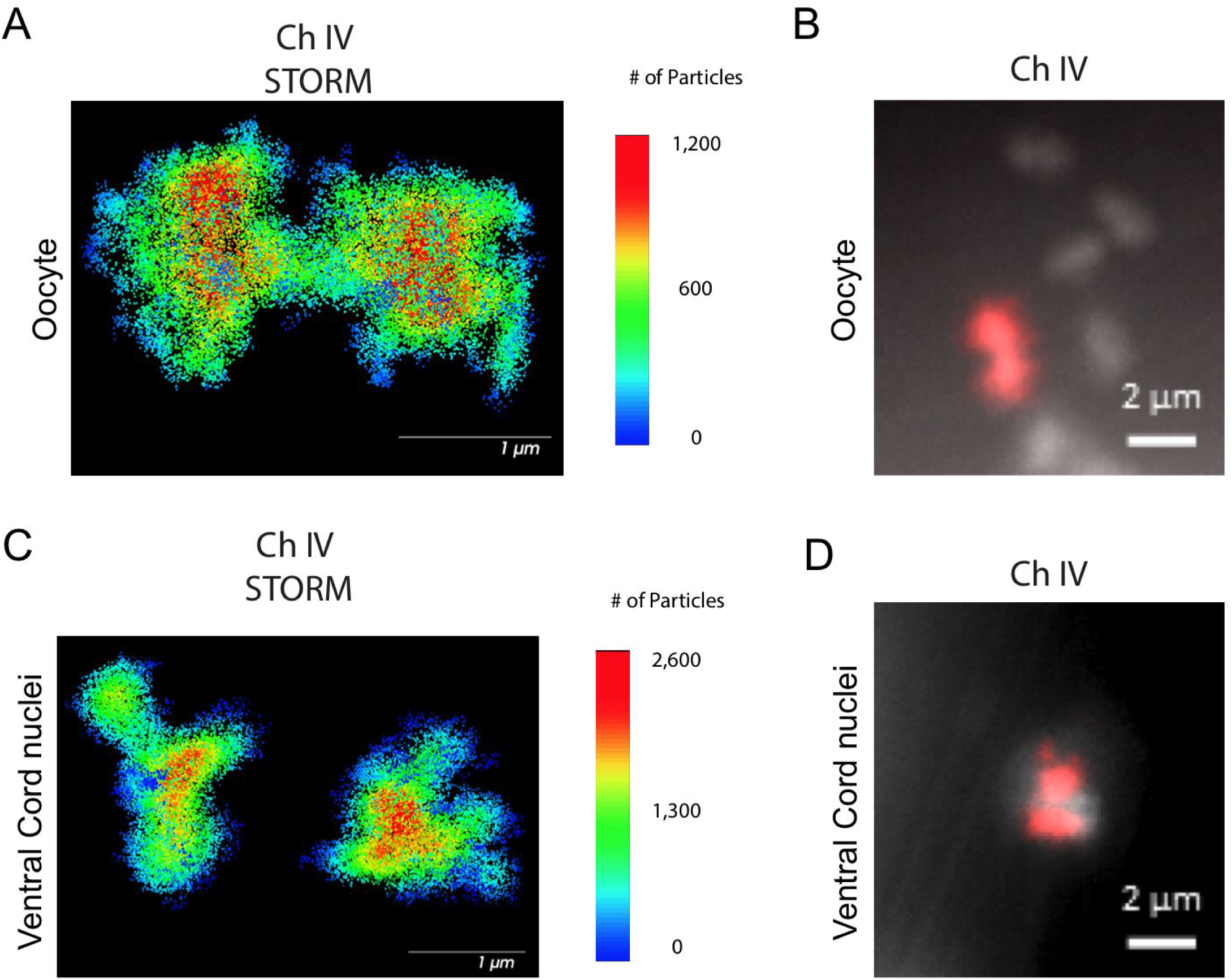
Coupling *C. elegans* Oligopaint to super resolution microscopy. Adult *C. elegans* were fixed and subjected to three step hybridization to detect Chromosome four with detection oligos suitable for STORM microscopy (Alexa647). **(A and C)** STORM microscopy of Alexa647 conjugated detection probes targeting chromosome IV in (**A)** an oocyte and **(C)** in a nucleus whose size and position within the animal suggests it is a ventral cord neuron. Data is displayed as particle density distributions using pair correlation analysis (see materials and methods). Scale bars are indicated. Fluorescent image using standard fluorescent microscope of an **(B)** oocyte nucleus and **(D)** a nucleus whose size and position within the animal suggests it is a ventral cord neuron.

### Using *C. elegans* Oligopaints to explore the biology of genome architecture

Our Oligopaints library and hybridization protocol will allow many questions relating to the biology of genome organization to be asked within the context of a whole animal. We started this process by using our library to ask two simple questions: 1) Whether genome architecture change during aging, and 2) what cellular factors are needed to establish and/or maintain chromosome territories in post-mitotic cells?

Recent studies suggest that higher order chromatin structures may break down during aging, and in aging related diseases such as Alzheimer’s (Winick-Ng and Rylett 2018). Age-related alterations in nuclear morphology have also been noted in *C. elegans* (Haithcock et al. 2005). We used our Oligopaint library to simultaneously visualize all 6 chromosomes in one and ten day old animals (*C. elegans* typically live about two weeks) to ask if the aging process might affect the compartmentalization of chromosomes into chromosome territories in *C. elegans.* We imaged intestinal nuclei as these cells are postmitotic, have large nuclei, and are easily identifiable due to their idiosyncratic size, shape, and location within the animal. As expected, all six chromosomes occupied largely distinct territories in intestinal cells of one day old animals. Interestingly, in ten day old animals, chromosomes were no longer organized into discrete territories (Fig. 6a). Quantification of the degree to which DNA FISH signals for Chromosomes I, II, and III overlapped in space (see materials and methods) supported the idea that chromosome territories are degraded in 10 day old nuclei (Fig. 6b). We asked if the loss of chromosome territories in older animals worms were a consequence of aging, or a function of time. To do so we conducted a similar analysis on young and old animals harboring a mutation (*e1370*) in *daf-2*. *daf-2* encodes a insulin-like receptor and loss-of-function mutations in *daf-2* cause animals to live twice as long as wild-type animals (Kenyon et al. 1993; Kimura et al. 1997). Chromosome territories were not enlarged or disorganized in young or ten day old *daf-2* mutant animals, indicating that the loss of chromosome territories we see in older wild-type animals is linked to aging and not chronological time (Fig. 6a/b). The data are consistent with the idea that higher-order chromatin structures are lost during the aging process in animals. Further studies will be needed to address if genome organization in other/all cell types is similarly affected by aging and, more importantly, if the loss of chromosomal territories is a cause or consequence of the aging process.

**Figure 6.**
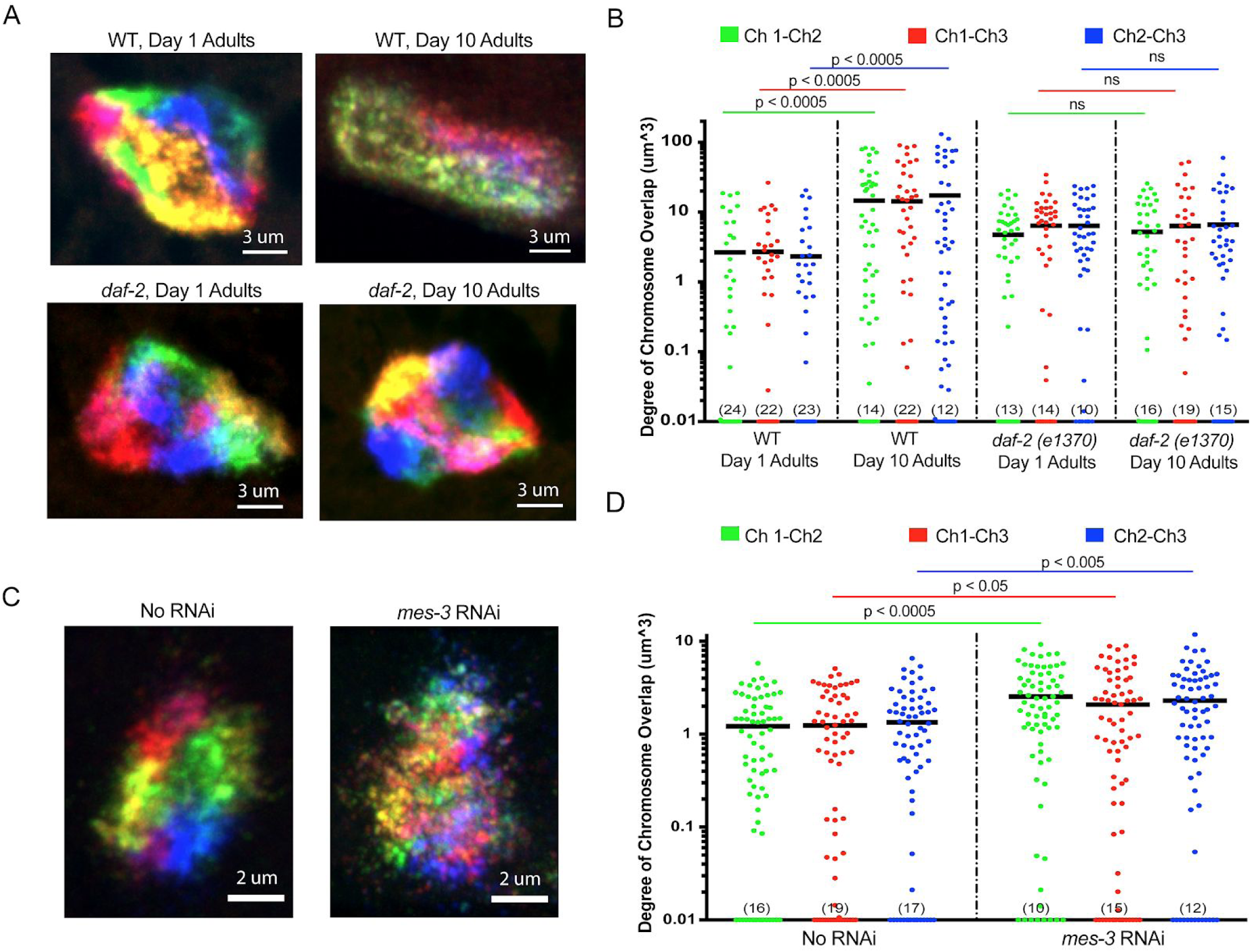
Using *C. elegans* Oligopaints to explore biology of genome organization in a whole animal. **(A)** Adult *C. elegans* were fixed and subjected to three step hybridization to detect all six chromosomes at day one or day ten of adulthood. 3D maximum projections of a representative intestinal nuclei is shown. Territories appear disorganized in ten day old animals. Chromosome territories in ten day old *daf-2(e1370)* (strain= CB1370) animals are not disorganized. **(B)** Experiment was repeated to detect just chromosomes I, II, and III (this was done to allow unambiguous identification of all three chromosomes-six color DNA FISH introduces overlapping fluorescence signals). ImageJ was used to quantify volume of overlap by chromosomes I, II, and III in one and ten day old animals (see materials and methods for details of quantification). We used volume of overlap as a metric of territory disorganization. The volume of overlap between chromosomes I and II, I and III, and II and III in one or ten day old animals of the indicated genotype is indicated on the y-axis. Numbers in parentheses indicate number of data points with value < 0.01. P values were calculated using a two-tailed student t-test. n = >50 nuclei from at least 5 animals **(C)**. Adult *C. elegans* were subjected to *mes-3* RNAi by feeding animals bacteria expressing *mes-3* dsRNA for two generations. Animals were subjected to three step hybridization to detect all six chromosomes and 3D maximum projections of a hypodermal nuclei is shown. Territories appear disorganized after treatment with *mes-3* dsRNA. **(D)** Quantifications of two independent three chromosome DNA FISH experiments reveals an increase in the volume of chromosome overlap after *mes-3* dsRNA. Numbers in parentheses indicate number of data points with value < 0.01. P values were calculated using a two-tailed student t-test. n = >50 nuclei from at least 5 animals.

Very little is known about how chromosome territories are established or maintained in animals. The Oligopaint DNA FISH library described above could be used to identify and characterize genes and pathways mediating these processes. As a first attempt to identify such factors, we conducted six chromosome DNA FISH on animals subjected to RNAi targeting seven candidate genes, which we suspected might be involved in establishing/maintaining chromosome territories in *C. elegans*. RNAi targeting the gene *mes-3*, a *C. elegans*-specific component of the Polycomb repressive complex 2 (PRC2), caused a loss of chromosome territories in adult hypodermal cells (Fig. 6c) (Xu et al. 2001; Capowski et al. 1991)(Holdeman et al. 1998; Ross and Zarkower 2003). Quantification of the degree to which DNA FISH signals for Chromosomes I, II, and III overlapped in wild-type animals and in animals subjected to mes-3 RNAi supported this conclusion (Fig. 6d). Note: we chose to image hypodermal cells for this analysis as these cells are, like intestinal cells, easy to identify, and because the effects of *mes-3* RNAi appeared, for unknown reasons, to be most dramatic in this cell type (data not shown). The data suggest that MES-3 and, therefore, PRC2 is needed to establish and/or maintain chromosome territories in *C. elegans*. Related studies using mutant alleles of *mes-3*, as well as loss-of-function alleles in other components of the PRC2, will be needed to confirm this link.

## Discussion

The invariant cell lineage, transparency, and small genome (100 Mb) of *C. elegans* make this animal an excellent system in which to explore how genome architecture relates to gene expression, development, growth, reproduction, and aging. DNA FISH experiments in *C. elegans* have historically relied on 1) labeling PCR products that cover a single small (5-10 kb) region, or 2) using BACs to generate probes targeting larger regions (up to a whole chromosomes). Such approaches are low throughput and rigid in the sense that new probe sets need to be produced for each new DNA FISH experiment. Such experiment have also been limited by the types of cells that can be visualized, as most DNA FISH protocols rely on dissection of tissues, which is not only low throughput, but also limits the number of cell types that can be analyzed at one time. Here we describe an Oligopaint DNA FISH library and hybridization strategy that, with a single PCR reaction, can generate probe sets that allow visualization of all six *C. elegans* chromosomes at varying scales. The ability to rapidly and cheaply produce *C. elegans* DNA FISH probes, in conjunction with improvements to hybridization protocols, enables DNA FISH in all/most cells across all stages of development in an intact animal. These improvements should empower studies asking if/how higher-order chromatin structures regulate, and/or are regulated by, changes in gene expression that occur during growth and development. Given the invariant cell lineage of *C. elegans*, it should now also be possible to ask if chromosome-chromosome interactions or homolog pairing, or the size, morphology, or sub-nuclear positioning, of chromosomal territories (or subregions of these territories) vary predictably by cell type, age, or developmental trajectory. The smallest unit of DNA detectable with our current library is 500kb. Future Oligopaint libraries could be designed to target smaller segments of DNA. For instance, we designed our library to have 2 probes/kb, meaning that each 500kb segment is targeted by ∼1000 primary probes. ∼200 have been shown to be sufficient for detection of Oligopaint signals, however (Beliveau et al. 2012). Additionally, our library includes only ∼20% of the high-quality sequences identified during our bioinformatic mining of the *C. elegans* genome. By incorporating all possible probes, and having each segment targeted by just 200 probes, future libraries could be designed to detect each and every 20 kb section of the *C. elegans* genome. Such a library could allow the architecture of the *C. elegans* genome to be visualized on an unprecedented scale, a process that might be even more powerful when coupled to super resolution microscopy, which we have shown is possible. The average *C. elegans* gene is about 3kb. Thus, Oligopaint libraries capable of individually detecting all ∼20,000 *C. elegans* genes are not yet possible. By targeting more fluorophores to each primary probe (using longer and more sophisticated bridge oligos), however, and by incorporating less-ideal (*e.g*. annealing temp) primary probes, single gene resolution might be achievable. Such a library could permit studies investigating the morphology, compaction state, or sub-nuclear distribution of any gene (or group of genes) within the context of a whole animal.

## Materials and Methods

### Design and synthesis of multiplexed DNA FISH library

We used a previously described pipeline to mine the *C. elegans* genome (build *ce10*) for desirable oligonucleotide nucleotide sequences 42 base pairs in length (Beliveau et al. 2012). 872,946 oligonucleotide sequences met this criteria, and 170,594 probes, which were evenly distributed across chromosomes, were chosen for the library. A series of barcode sequences were appended to each 42 bp hybridization sequence, which resulted in each primary probe being 150 bp. Barcode sequences can be found in Supplemental table 2. The 170,594 sequences were ordered as two 90k oligonucleotide chips from Custom Array (Bothell, WA). To obtain primary probes for Oligopaint experiments, desired primary probes were first amplified using primers specific to the outermost barcode sequences, which correspond to the individual chromosome barcodes shown in Figure 1a. Single stranded probe (primary probe) generation was conducted as previously described (Chen et al. 2015). Briefly, PCR was used to append a T7 polymerase site to the 5’ end of chromosome specific barcode sequence, followed by T7 polymerase reactions to generate ssRNA. ssRNA was reverse transcribed into ssDNA. Unwanted ssRNA species were degraded using base hydrolysis. Finally, long ssDNA oligos were purified using the Zymo-100 DNA Clean and Concentrator Kit with oligo binding buffer. Probes were stored at 100 pmol/ul at -20C.

### Bridge and Detection probes

Bridge oligos were ordered from IDT at 25 or 100 nmole scales using standard desalting procedures. Fluorescent detection oligos were ordered from IDT with 5’ and 3’ fluorescent modifications on the 250 nm or 1 um scale and subjected to HPLC purification. Bridge and detection probe sequences are listed in Supplemental table 2.

### DNA FISH

Embryos were isolated by hypochlorite treatment and fed OP50 bacteria seeded onto 10 cm agar plates. When animals reached adulthood, plates were washed with M9 solution and collected in 15 ml conical tubes. Animals were pelleted (3k rpm for 30 seconds), and washed 2 times with M9 solution. Animals were resuspended in 10 ml of M9 solution and rocked for ∼30 min at room temperature. Animals were pelleted and aliquoted to 1.5 ml microcentrifuge tubes (50 ul of packed worms per tube). Samples were placed in liquid nitrogen for 1 minute. For mixed stage analysis, populations of mixed stage worms were transferred to 10 cm plates. Prior to starvation, animals were washed off the plate with M9 buffer and transferred to 15 ml conical tubes. Animals were washed twice with M9 solution and rocked for ∼30 min at room temperature. Animals were pelleted and aliquoted to 1.5 ml microcentrifuge tubes (50 ul of packed worms per tube). Samples were placed in liquid nitrogen for 1 minute. Frozen worm pellets were resuspended in cold 95% ethanol and vortexed for 30 seconds. Samples were rocked for 10 minutes at room temperature. Samples were spun down (3k rpm for 30 seconds), supernatant discarded, and washed twice in 1X PBST. 1 ml of 4% paraformaldehyde solution was added and samples were rocked at room temperature for 5 minutes, washed twice with 1X PBST, and resuspended in 2XSSC for 5 minutes at room temperature. Samples were spun down and resuspended in a 50% formamide 2XSSC solution at room temperature for 5 minutes, 95°C for 3 minutes, and 60°C for 20 minutes. Samples were spun and resuspended in 60 ul of hybridization mixture (10% dextran sulfate, 2XSSC, 50% formamide, 100 pmol of primary probe per chromosome and 2 ul of RNAse A (sigma 20 mg/ml)). Hybridization reactions were incubated in a 100°C heat block for 5 minutes before overnight incubation at 37°C in a hybridization oven. The next day, samples were washed with prewarmed 2XSSCT (rotating at 60°C) for 5 minutes, followed by a second 2XSSCT wash at 60°C for 20 minutes. Wash buffer was removed and samples were resuspended in 60 ul of bridge oligo hybridization mixture (2XSSC, 30% formamide, 100 pmol of bridge oligo per targeted region (ie whole chromosome, three megabase, or 500 kb spots) and 100 pmol of each detection oligo. Bridge/detection oligo hybridization reactions were incubated at room temperature for 3 hours. Samples were washed in prewarmed 2XSSC at 60°C for 20 minutes, followed by a 5 minute wash with 2XSSCT at 60°C and a 20 minute wash in 2XSSCT at 60°C. Samples were then washed at room temperature in 2XSSCT. Wash buffer was removed and samples were resuspended in mounting medium (vectashield with DAPI or slowfade Gold with DAPI if imaging Alexa647--Alexa647 is not photostable in vectashield). Samples were mounted on microscope slides and sealed with nail polish.

### Alternate embryo DNA FISH protocol

DNA FISH on in utero embryos was only 50% efficient. The following protocol improves this efficiency to nearly 100%. This protocol is an adaptation of an existing *C. elegans* DNA FISH protocol (Crane et al. 2015). Briefly, adults were dissected in 8 ul of 1X egg buffer on a coverslip (25 mM HEPEs, pH 7.3, 118 mM NaCl_2_, 48 mM KCl, 2 mM CaCl_2_, 2 mM MgCl_2_) to release embryos. Coverslips were placed on a Superfrost Plus Gold slide (Thermo Scientific) and placed in liquid nitrogen for one minute. Coverslips were popped off with a razor blade and slides were submerged in 95% cold (−20C) ethanol for 10 minutes. Slides were washed twice in 1XPBST before fixation in 4% paraformaldehyde solution (described above) for 5 minutes. Slides were washed twice in 1XPBST. 20 ul Primary hybridization mixture (described above) was added to each sample and a coverslip was placed on top. Slides were placed on a 90°C heat block for 10 minutes. Slides were placed in a humid chamber at 37°C overnight. Wash steps and secondary/bridge hybridization was carried out as described above. 15 ul of mounting medium was added to each sample, and coverslips were sealed with nail polish.

### Microscopy

Standard fluorescent microscopy was conducted on a widefield Zeiss Axio Observer.Z1 microscope using a Plan-Apochromat 63X/1.40 Oil DIC M27 objective and an ORCA-Flash 4.0 CMOS Camera. The Zeiss Apotome 2.0 was used for structured illumination microscopy using 3 phase images. All image processing was done using the Zen imaging software from Zeiss. Confocal microscopy was done using a Nikon Eclipse Ti microscope equipped with a W1 Yokogawa Spinning disk with 50 um pinhole disk and an Andor Zyla 4.2 Plus sCMOS monochrome camera. A 60X/1.4 Plan Apo Oil objective was use.

### STORM

STORM imaging was performed on Vutara 352 (Bruker) commercial 3D biplane single molecule localization microscope, equipped with 60X water objective (Olympus) with a numerical aperture (NA) of 1.2. For sample illumination, we used 640-nm laser with ∼100 mW at the back aperture of the objective; for imaging fluorescent beads, as fiducial drift-correction, illumination at the back aperture of the objective was ∼3 mW for the 488-nm laser. Fluorescent detection was captured on an sCMOS camera (Hamamatsu Orca-flash 4.0 v2) using a 10 ms exposure time. We collected > 75,000 frames for each locus, while taking up to 5 μm z-scan using 100 nm z-steps.

### STORM data analysis

Drift correction was performed using the fiducial alignment function within Vutara software. Six fiducial beads were used for each alignment. STORM data was denoised using the Vutara software prior to resolution analysis. Cluster analysis was performed using DBscan within the Vutara software, and particle density distributions were calculated using pair correlation analysis within the Vutara software.

### Assessment of overlapping chromosome territory volumes

All territory quantifications were done using standard tools in ImageJ along with the 3D objects counter plugin (Bolte and Cordelières 2006). First, each individual nuclei was segmented from the original file to generate individual nuclei files. The 4 channel stack was then split to create individual files for each chromosome (each chromosome is represented by a single fluorophore/channel). To remove background noise and create a binary mask, each image was subjected to thresholding using the default ImageJ thresholding using “auto” across every image. Once masks were obtained the 3D objects counter tool was utilized to select objects larger than 30 voxels (eliminating further background signal). Object masks for each channel were loading into the 3D Manager plugin for ImageJ, and all objects for a given chromosome were merged into a single object. The colocalization function within 3D manager was used to determine the volume of overlap between each chromosome.

## Acknowledgments

We would like to thank members of the Kennedy and Wu Labs for helpful discussions. We also thank Barbara Meyer and Satoru Uzawa for helpful discussions on DNA FISH in *C. elegans* embryos. BF and SN were supported by NSF graduate research fellowships. This work was supported by the National Institutes of Health, RO1 GM088289 (S.K.)

## Figures

**Movie 1. Six Chromosome FISH of a *C. elegans* intestinal nucleus.** 3D render of a single *C. elegans* intestinal nucleus from a worm hybridized with the 6 chromosome Oligopaint strategy. Individual chromosome territories can be clearly distinguished.

**Movie 2. Chromosome 4 STORM of a *C. elegans* oocyte.** 3D rotation of the oocyte depicted in Fig 5a. Data is displayed as particle density distributions using pair correlation analysis. Note that *C. elegans* oocytes are stalled in diakinesis, where each homologous chromosome remains attached via a chiasmata. Each homologous chromosome, as well as the chiasmata can be clearly distinguished. Particle density profiles reveal core like structures at the center of each homolog.

## Supplementary Figures and Tables

**Supplementary File 1. Barcode, bridge, and detection oligo sequences used in this study.** This file contains the barcodes, bridge, and detection oligos used in this study, as well as the fluorophores used to label each detection oligo. Similar bridge and detection oligos could in theory be used for any primary probe targeting any genome.

**Supplementary Movie 1. Chromosome IV STORM of a *C. elegans* oocyte--Z slices.** Z slice representation of the oocyte depicted in Fig 6a and Movie 2. Individual Z slices are shown to allow visualization of individual particles. Data is displayed as particle density distributions using pair correlation analysis.

**Supplementary Movie 2. Chromosome IV STORM of a *C. elegans* ventral cord neuron--Z slices.** Z slice representation of the ventral cord neuron depicted in Fig 6b. Individual Z slices are shown to allow visualization of individual particles. Data is displayed as particle density distributions using pair correlation analysis.

**Supplementary Movie 3. Chromosome IV STORM of a *C. elegans* ventral cord neuron--3D rotation.** 3D rotation of the oocyte depicted in Fig 6b. Data is displayed as particle density distributions using pair correlation analysis. Note that like the C. elegans oocytes, the individual homologous chromosomes display core-like structures in the center of the chromosome territory in the interphase neuronal nuclei.

**Supplementary Movie 4. Chromosome III STORM of a *C. elegans* oocyte--Z slices.** Z slice representation of the oocyte depicted in Sup Fig 1a. Individual Z slices are shown to allow visualization of individual particles. Data is displayed as particle density distributions using pair correlation analysis.

**Supplementary Movie 5. Chromosome III STORM of a *C. elegans* oocyte--3D rotation.** 3D rotation of the oocyte depicted in Sup Fig 1a. Data is displayed as particle density distributions using pair correlation analysis. Note that like Chromosome IV, Chromosome III also displays dense signal in the center of each homologous chromosome.

**Supplementary Movie 6. Chromosome III STORM of a *C. elegans* intestinal nucleus--Z slices.** Z slice representation of the intestinal nucleus depicted in Sup Fig 1b. Individual Z slices are shown to allow visualization of individual particles. Data is displayed as particle density distributions using pair correlation analysis.

**Supplementary Movie 7. Chromosome III STORM of a *C. elegans* oocyte--3D rotation**. 3D rotation of the intestinal nucleus depicted in Sup Fig 1b. Data is displayed as particle density distributions using pair correlation analysis. Core-like structures can also be observed in the chromosome territories of interphase intestinal nuclei.

**Figure S1.**
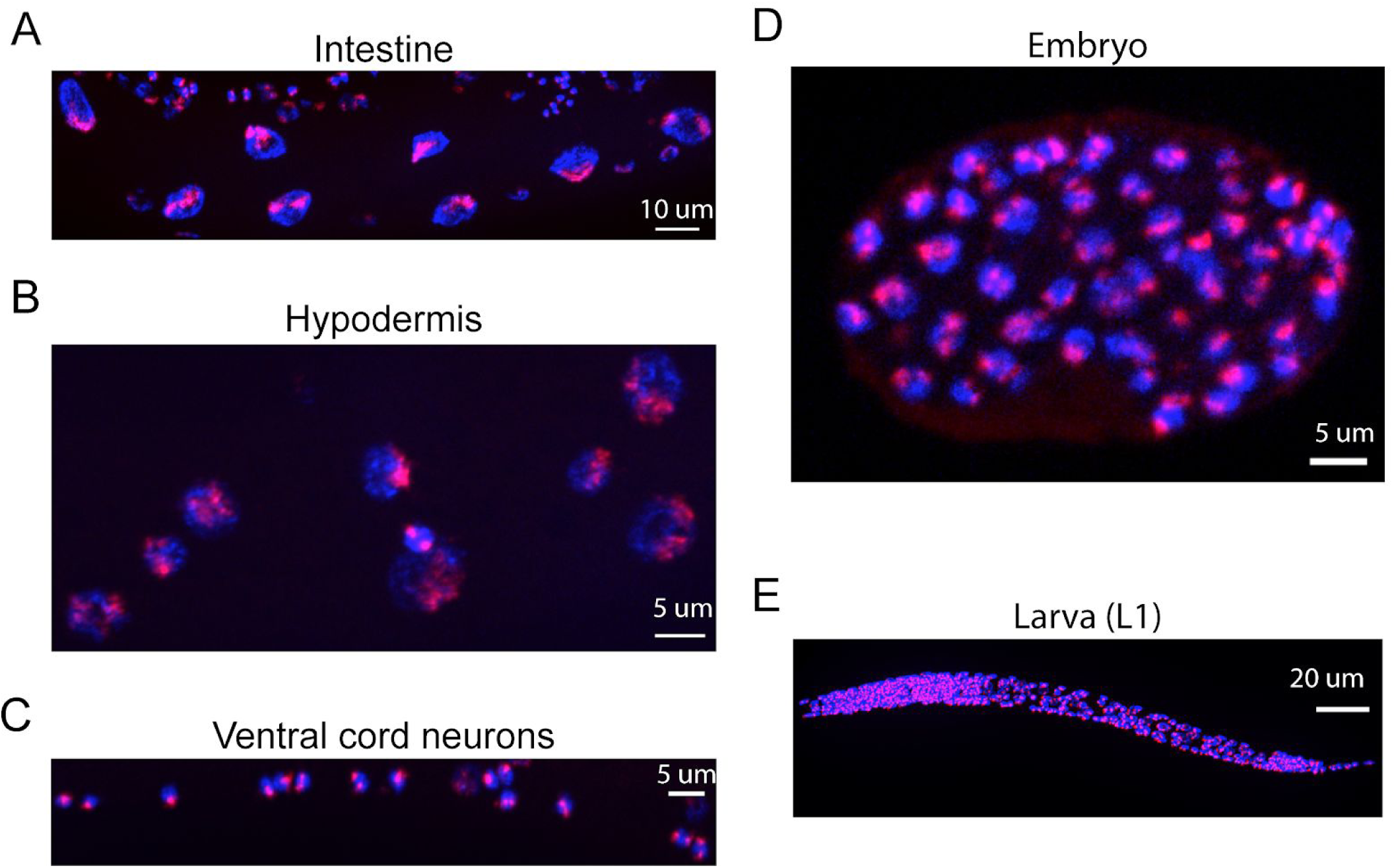
Oligopaints stain chromosomes in all/most somatic cells and in all/most developmental stages of *C. elegans.* Mixed stage samples of C. elegans were subjected to three step hybridization to detect Chromosome II as described in materials and methods. Chromosome II staining is shown in red. Animals were co-stained with DAPI (blue). Scale bars are indicated. **(A)** Intestinal nuclei, **(B)** hypodermal nuclei, **(C)** ventral cord neurons, **(D)** ∼100 cell embryo, and **(E)** larval stage one animal.

**Figure S2.**
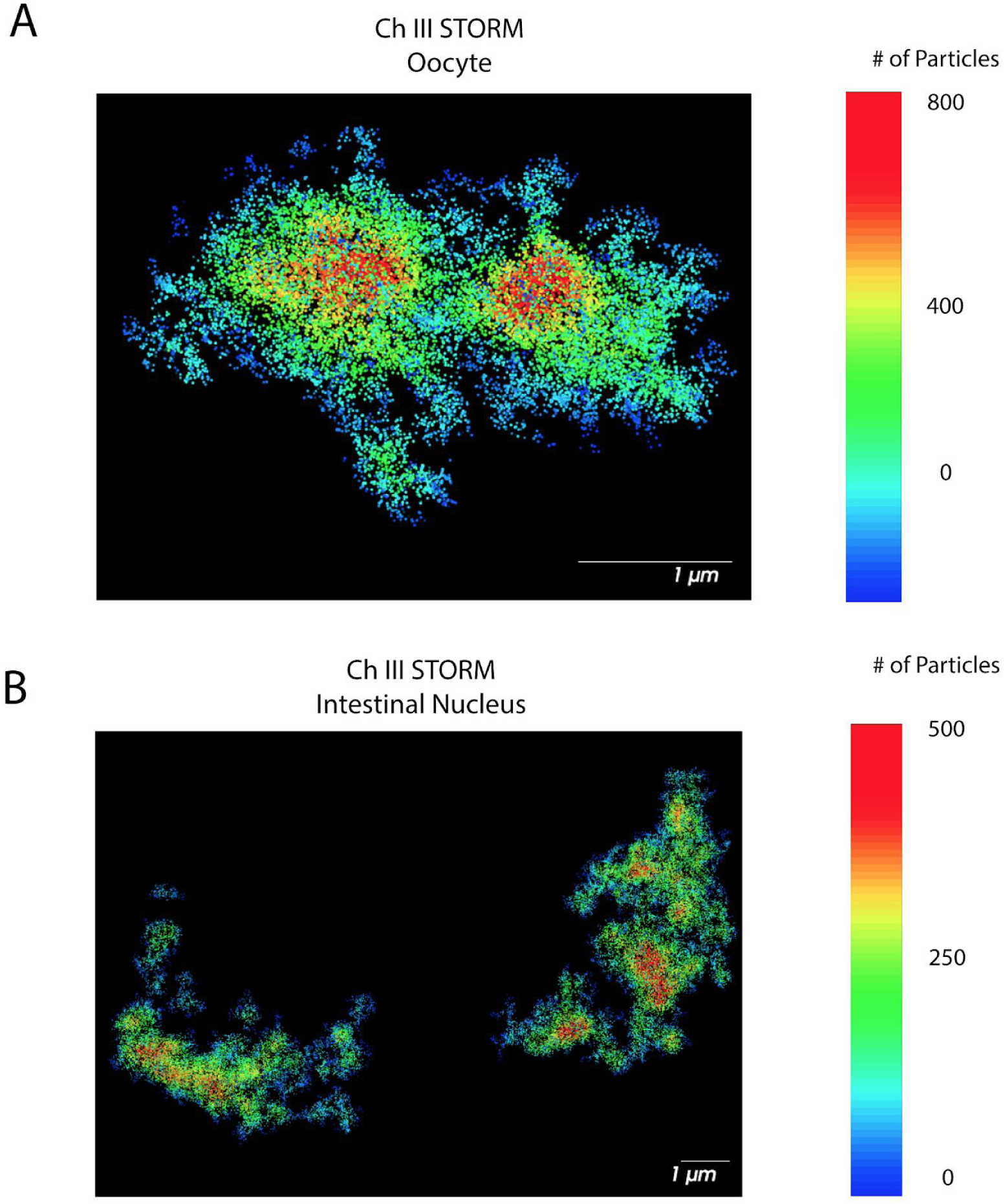
STORM of *C. elegans* Chromosome III in an oocyte and intestinal nucleus. Adult C. elegans were fixed and subjected to three step hybridization to detect chromosome three with a detection oligo suitable for STORM microscopy (Alexa647). **(A-B)** STORM microscopy of chromosome three of an oocyte **(A)** and intestinal nucleus **(B)**. Data is displayed as particle density distributions using pair correlation analysis (see materials and methods). Scale bars are indicated. Image shows a densely packed region of DNA in the interior of chromosome three.

**Supplementary Table 1.** Distribution of oligonucleotide sequences across the *C. elegans* genome. Potential probes sequences were identified as described in materials and methods. Every 5th oligo sequence was incorporated into library to ensure even distribution. The average distance between selected probes for each chromosome, and the standard deviations of these distances, are indicated. Kb, kilobase. bp, base pair.

| Chr | First Coordinate | Last Coordinate | Potential Oligo Sequences | Potential Oligo Sequences/kb | Oligo Sequences Chosen | Oligo Sequences Chosen/kb | Average distance between oligos (bp) | Standard Deviation of distance between oligos (bp) |
| --- | --- | --- | --- | --- | --- | --- | --- | --- |
| I | 512 | 14999984 | 125,863 | 8.39 | 25,174 | 1.68 | 595.8 | 505.9 |
| II | 163 | 15199262 | 136,793 | 9.00 | 27,360 | 1.80 | 555.5 | 484.5 |
| III | 123 | 13599784 | 133,977 | 9.85 | 22,796 | 1.68 | 595.9 | 572.2 |
| IV | 655 | 17399857 | 137,270 | 7.89 | 27,454 | 1.58 | 633.8 | 585.4 |
| V | 414 | 20699907 | 177,948 | 8.60 | 35,590 | 1.72 | 581.6 | 460.4 |
| X | 595 | 17599761 | 161,095 | 9.15 | 32,220 | 1.83 | 546.2 | 436.8 |
|  |  |  | 872,946 | 8.81 | 170,594 | 1.71 | 584.8 | 507.5 |

## Bibliography

Albertson, Donna G., Ann M. Rose, and Anne M. Villeneuve. 2011. “Chromosome Organization, Mitosis, and Meiosis.” In C. Elegans II, edited by Donald L. Riddle, Thomas Blumenthal, Barbara J. Meyer, and James R. Priess. Cold Spring Harbor (NY): Cold Spring Harbor Laboratory Press.

Banterle, Niccolò, Khanh Huy Bui, Edward A. Lemke, and Martin Beck. 2013. “Fourier Ring Correlation as a Resolution Criterion for Super-Resolution Microscopy.” Journal of Structural Biology 183 (3): 363–67.

Bauman, J. G., J. Wiegant, P. Borst, and P. van Duijn. 1980. “A New Method for Fluorescence Microscopical Localization of Specific DNA Sequences by in Situ Hybridization of Fluorochromelabelled RNA.” Experimental Cell Research 128 (2): 485–90.

Beliveau, Brian J., Alistair N. Boettiger, Maier S. Avendaño, Ralf Jungmann, Ruth B. McCole, Eric F. Joyce, Caroline Kim-Kiselak, et al. 2015. “Single-Molecule Super-Resolution Imaging of Chromosomes and in Situ Haplotype Visualization Using Oligopaint FISH Probes.” Nature Communications 6 (May): 7147.

Beliveau, Brian J., Eric F. Joyce, Nicholas Apostolopoulos, Feyza Yilmaz, Chamith Y. Fonseka, Ruth B. McCole, Yiming Chang, et al. 2012. “Versatile Design and Synthesis Platform for Visualizing Genomes with Oligopaint FISH Probes.” Proceedings of the National Academy of Sciences 109 (52): 21301–6.

Bickmore, Wendy A. 2013. “The Spatial Organization of the Human Genome.” Annual Review of Genomics and Human Genetics 14 (1): 67–84.

Bienko, Magda, Nicola Crosetto, Leonid Teytelman, Sandy Klemm, Shalev Itzkovitz, and Alexander van Oudenaarden. 2013. “A Versatile Genome-Scale PCR-Based Pipeline for High-Definition DNA FISH.” Nature Methods 10 (2): 122–24.

Boettiger, Alistair N., Bogdan Bintu, Jeffrey R. Moffitt, Siyuan Wang, Brian J. Beliveau, Geoffrey Fudenberg, Maxim Imakaev, Leonid A. Mirny, Chao-Ting Wu, and Xiaowei Zhuang. 2016. “Super-Resolution Imaging Reveals Distinct Chromatin Folding for Different Epigenetic States.” Nature 529 (7586): 418–22.

Bolte, S., and F. P. Cordelières. 2006. “A Guided Tour into Subcellular Colocalization Analysis in Light Microscopy.” Journal of Microscopy 224 (Pt 3): 213–32.

Bolzer, Andreas, Gregor Kreth, Irina Solovei, Daniela Koehler, Kaan Saracoglu, Christine Fauth, Stefan Müller, et al. 2005. “Three-Dimensional Maps of All Chromosomes in Human Male Fibroblast Nuclei and Prometaphase Rosettes.” PLoS Biology 3 (5): e157.

Bonev, Boyan, and Giacomo Cavalli. 2016. “Organization and Function of the 3D Genome.” Nature Reviews. Genetics 17 (11): 661–78.

Casolari, Jason M., Christopher R. Brown, Suzanne Komili, Jason West, Haley Hieronymus, and Pamela A. Silver. 2004. “Genome-Wide Localization of the Nuclear Transport Machinery Couples Transcriptional Status and Nuclear Organization.” Cell 117 (4): 427–39.

Chen, Kok Hao, Alistair N. Boettiger, Jeffrey R. Moffitt, Siyuan Wang, and Xiaowei Zhuang. 2015. “RNA Imaging. Spatially Resolved, Highly Multiplexed RNA Profiling in Single Cells.” Science 348 (6233): aaa6090.

Crane, Emily, Qian Bian, Rachel Patton McCord, Bryan R. Lajoie, Bayly S. Wheeler, Edward J. Ralston, Satoru Uzawa, Job Dekker, and Barbara J. Meyer. 2015. “Condensin-Driven Remodelling of X Chromosome Topology during Dosage Compensation.” Nature 523 (7559): 240–44.

Cremer, Thomas, and Marion Cremer. 2010. “Chromosome Territories.” Cold Spring Harbor Perspectives in Biology 2 (3). https://doi.org/10.1101/cshperspect.a003889.

Dekker, Job, and Edith Heard. 2015. “Structural and Functional Diversity of Topologically Associating Domains.” FEBS Letters 589 (20 Pt A): 2877–84.

Dekker, Job, Marc A. Marti-Renom, and Leonid A. Mirny. 2013. “Exploring the Three-Dimensional Organization of Genomes: Interpreting Chromatin Interaction Data.” Nature Reviews. Genetics 14 (6): 390–403.

Dekker, Job, and Leonid Mirny. 2016. “The 3D Genome as Moderator of Chromosomal Communication.” Cell 164 (6): 1110–21.

Dekker, Job, Karsten Rippe, Martijn Dekker, and Nancy Kleckner. 2002. “Capturing Chromosome Conformation.” Science 295 (5558): 1306–11.

Dixon, Jesse R., Siddarth Selvaraj, Feng Yue, Audrey Kim, Yan Li, Yin Shen, Ming Hu, Jun S. Liu, and Bing Ren. 2012. “Topological Domains in Mammalian Genomes Identified by Analysis of Chromatin Interactions.” Nature 485 (7398): 376–80.

Gonzalez-Sandoval, Adriana, and Susan M. Gasser. 2016. “On TADs and LADs: Spatial Control Over Gene Expression.” Trends in Genetics: TIG 32 (8): 485–95.

Haithcock, Erin, Yaron Dayani, Ester Neufeld, Adam J. Zahand, Naomi Feinstein, Anna Mattout, Yosef Gruenbaum, and Jun Liu. 2005. “Age-Related Changes of Nuclear Architecture in Caenorhabditis Elegans.” Proceedings of the National Academy of Sciences of the United States of America 102 (46): 16690–95.

Kenyon, C., J. Chang, E. Gensch, A. Rudner, and R. Tabtiang. 1993. “A C. Elegans Mutant That Lives Twice as Long as Wild Type.” Nature 366 (6454): 461–64.

Kimura, K. D., H. A. Tissenbaum, Y. Liu, and G. Ruvkun. 1997. “Daf-2, an Insulin Receptor-like Gene That Regulates Longevity and Diapause in Caenorhabditis Elegans.” Science 277 (5328): 942–46.

Lemaître, Charlene, and Wendy A. Bickmore. 2015. “Chromatin at the Nuclear Periphery and the Regulation of Genome Functions.” Histochemistry and Cell Biology 144 (2): 111–22.

Lieberman-Aiden, Erez, Nynke L. van Berkum, Louise Williams, Maxim Imakaev, Tobias Ragoczy, Agnes Telling, Ido Amit, et al. 2009. “Comprehensive Mapping of Long-Range Interactions Reveals Folding Principles of the Human Genome.” Science 326 (5950): 289–93.

Meaburn, Karen J. 2016. “Spatial Genome Organization and Its Emerging Role as a Potential Diagnosis Tool.” Frontiers in Genetics 7 (July): 134.

Murgha, Yusuf E., Jean-Marie Rouillard, and Erdogan Gulari. 2014. “Methods for the Preparation of Large Quantities of Complex Single-Stranded Oligonucleotide Libraries.” PloS One 9 (4): e94752.

Pickersgill, Helen, Bernike Kalverda, Elzo de Wit, Wendy Talhout, Maarten Fornerod, and Bas van Steensel. 2006. “Characterization of the Drosophila Melanogaster Genome at the Nuclear Lamina.” Nature Genetics 38 (9): 1005–14.

Rust, Michael J., Mark Bates, and Xiaowei Zhuang. 2006. “Sub-Diffraction-Limit Imaging by Stochastic Optical Reconstruction Microscopy (STORM).” Nature Methods 3 (10): 793–96.

Steensel, Bas van, and Andrew S. Belmont. 2017. “Lamina-Associated Domains: Links with Chromosome Architecture, Heterochromatin, and Gene Repression.” Cell 169 (5): 780–91.

Vernimmen, Douglas, and Wendy A. Bickmore. 2015. “The Hierarchy of Transcriptional Activation: From Enhancer to Promoter.” Trends in Genetics: TIG 31 (12): 696–708.

Villeneuve, A. M. 1994. “A Cis-Acting Locus That Promotes Crossing over between X Chromosomes in Caenorhabditis Elegans.” Genetics 136 (3): 887–902.

Winick-Ng, Warren, and R. Jane Rylett. 2018. “Into the Fourth Dimension: Dysregulation of Genome Architecture in Aging and Alzheimer’s Disease.” Frontiers in Molecular Neuroscience 11 (February): 60.

Yu, Miao, and Bing Ren. 2017. “The Three-Dimensional Organization of Mammalian Genomes.” Annual Review of Cell and Developmental Biology 33 (October): 265–89.

